# Neural distinctiveness declines with age in auditory cortex and is associated with auditory GABA levels

**DOI:** 10.1101/470781

**Authors:** Poortata Lalwani, Holly Gagnon, Kaitlin Cassady, Molly Simmonite, Scott Peltier, Rachael D. Seidler, Stephan F. Taylor, Daniel H. Weissman, Thad A. Polk

**Author notes:** Corresponding author Thad Polk, University of Michigan, Department of Psychology, 530 Church Street, Ann Arbor, MI 48109, USA.

## Abstract

Neural activation patterns in the ventral visual cortex in response to different categories of visual stimuli (e.g., faces vs. houses) are less selective, or distinctive, in older adults than in younger adults, a phenomenon known as age-related neural dedifferentiation. Previous work in animals suggests that age-related reductions of the inhibitory neurotransmitter, gamma aminobutyric acid (GABA), may play a role in this age-related decline in neural distinctiveness. In this study, we investigated whether neural dedifferentiation extends to auditory cortex and whether individual differences in GABA are associated with individual differences in neural distinctiveness in humans. 20 healthy young adults (ages 18-29) and 23 healthy older adults (over 65) completed a functional magnetic resonance imaging (fMRI) scan, during which neural activity was estimated while they listened to foreign speech and music. GABA levels in the auditory, ventrovisual and sensorimotor cortex were estimated in the same individuals in a separate magnetic resonance spectroscopy (MRS) scan. Relative to the younger adults, the older adults exhibited both (1) less distinct activation patterns for music vs. speech stimuli and (2) lower GABA levels in the auditory cortex. Also, individual differences in auditory GABA levels (but not ventrovisual or sensorimotor GABA levels) predicted individual differences in neural distinctiveness in the auditory cortex in the older adults. These results demonstrate that age-related neural dedifferentiation extends to the auditory cortex and suggest that declining GABA levels may play a role in neural dedifferentiation in older adults.

**Significance Statement:** Prior work has revealed age-related neural dedifferentiation in the visual cortex. GABA levels also decline with age in several parts of the human cortex. Here, we report that these two age-related changes are linked; neural dedifferentiation is associated with lower GABA levels in older adults. We also show that age-related neural dedifferentiation extends to auditory cortex, suggesting that it may be a general feature of the aging brain. These findings provide novel insights into the neurochemical basis of age-related neural dedifferentiation in humans and also offer a potential new avenue for investigating age-related declines in central auditory processing.

## Introduction

Aging is often accompanied by declines in cognitive (1–3) and sensory (4) function. These declines have a significant negative impact on the daily lives of older individuals and are often early indicators of pathology. However, there are significant individual differences in these declines: some older adults experience severe impairments while others do not (5–7). Understanding the neural bases of these individual differences may therefore be helpful in designing interventions that slow or halt some age-related impairments.

One neural factor that may play a role is an age-related decline in neural distinctiveness. Neural activity associated with different categories of visual stimuli (e.g., faces, houses, words) is often less selective, or distinctive, in older adults than in younger adults (8–10). Individual differences in this kind of age-related “neural dedifferentiation” are also a significant predictor of cognitive decline. For example, older adults with less distinctive neural activation patterns perform more poorly on a wide range of fluid processing tasks compared with those with more distinctive patterns (11).

Another neural factor that may be important is age-related reductions in the brain’s major inhibitory neurotransmitter, gamma-aminobutyric acid (GABA). GABA levels measured using magnetic resonance spectroscopy (MRS) have been found to be reduced in older adults compared to younger adults in the occipital cortex (12–14), in frontal and parietal regions (13, 15), and in supplementary motor area and sensorimotor cortex (12, 13, 16). Furthermore, individual differences in GABA in specific cortical regions have been associated with individual differences in some aspects of cognitive performance (13, 14, 17).

To date, age-related neural dedifferentiation and declines in GABA levels have been studied in isolation from one another. In the present study, we test whether individual differences in GABA predict individual differences in neural distinctiveness and if this relationship is region specific. This work is motivated by previous studies in animals showing a causal link between GABA levels and neural selectivity. Leventhal et al. (2003) showed that the application of GABA or a GABA agonist increased the orientation selectivity of cells in the visual cortex of older rhesus monkeys. Conversely, application of a GABA antagonist decreased the orientation selectivity of cells in the visual cortex of young monkeys (18). GABA receptor antagonists have also been shown to broaden the frequency response of neurons in the inferior colliculus of chinchillas, making the cells’ response less selective (19).

Inspired by these findings, we investigate the relationship between GABA levels and neural distinctiveness in the human auditory cortex. Most previous neural dedifferentiation studies have been focused on the visual cortex, and so it remains unclear whether dedifferentiation also occurs in other sensory regions, such as the auditory cortex. Furthermore, the results of the small number of human studies investigating age-related changes in GABA levels in the auditory cortex are mixed (20, 21). In the present study, we therefore asked whether older adults would have reduced distinctiveness and reduced GABA levels in the auditory cortex compared with young adults.

Research in animals suggests that the answer may be yes. For example, Turner, Hughes, & Caspary (2005) reported that the receptive fields of auditory neurons are less selective to pure tones in older rats compared with younger rats. Frequency selective bandwidths of auditory neurons get larger and receptive fields overlap more in older rats (22). Likewise, neurons in primary and secondary auditory cortex are less spatially tuned in older compared with younger macaques (23). Auditory frequency selectivity also declines with age in mice (24). Together, these results suggest that in many mammals, neural selectivity declines not only in visual cortex but also in auditory cortex. In this study, we used functional MRI to examine whether activation patterns for different categories of auditory stimuli are less distinct in human beings as well.

Studies have also reported age-related decreases in the protein and mRNA levels of the most abundant GABA_A_ receptor subunits in inferior colliculus and auditory cortex of rats (25–27). GABA_B_ receptor binding in the inferior colliculus also declines with age in rats (28). In this study, we used Magnetic Resonance Spectroscopy (MRS) to examine whether GABA levels measured in the human auditory cortex decline with age. We also examined whether individual differences in GABA are associated with individual differences in neural distinctiveness.

In sum, we combined fMRI and MRS to test whether age-related dedifferentiation extends to the human auditory cortex, whether auditory GABA levels decline with age, and whether GABA levels and neural distinctiveness are associated in the auditory cortex of older adults.

## Methods

### Participants

Twenty young adults (8 males, mean age = 23.6, range 18 to 28 years) and 23 older adults (7 males, mean age = 69.91, range 65 to 81 years) adults participated in the study. All participants were right-handed, native English speakers with normal or corrected to normal vision. We excluded participants who used hearing aids or scored lower than 23 on the Montreal Cognitive Assessment (MOCA) (29). We ensured that none of our participants knew any of the foreign languages that were used as auditory stimuli for the fMRI task. All sessions took place at the University of Michigan’s Functional MRI Laboratory, Ann Arbor, Michigan. Participants were recruited from Ann Arbor and the surrounding area.

### Session Design

Eligible participants completed a functional MRI session and an MRS session on the same scanner on separate days within a few weeks of each other. These data were collected as a part of larger study called the Michigan Neural Distinctiveness or MiND study. Here, we only describe the portions of the study that are relevant to this experiment. Please refer to (30) for further details on the MiND study itself.

### fMRI Session

We collected both structural and functional MRI data using a 3T General Electric Discovery Magnetic Resonance System with an 8-channel head coil at the Functional MRI Laboratory, University of Michigan, Ann Arbor, MI, USA. We obtained T1-weighted images using an SPGR (3D BRAVO) sequence with the following parameters: Inversion Time (TI) = 500 ms; flip angle = 15°; Field of View (FOV) = 256 x 256 mm. While the structural scan was being collected, each participant heard a trial version of the auditory stimuli and the volume was adjusted to ensure that each participant could comfortably hear the stimuli presented during the scan.

During the functional scans, T2*-weighted images were collected with a 2D Gradient Echo spiral pulse sequence with the following parameters: TR = 2000 ms; TE = 30 ms; flip angle = 90°; FOV = 220 x 220 mm; 43 axial slices with thickness = 3 mm and no spacing, collected in an interleaved bottom-up sequence. The total acquisition time for the functional scan was 6 minutes and 10 seconds with 185 volumes. E-Prime software was used to present auditory stimuli, which consisted of six 20-second blocks of foreign speech clips, six 20-second blocks of instrumental music clips, and twelve 10-second blocks of fixation between every pair of auditory blocks. The order of the speech and music blocks was pseudorandomized.

Each speech block consisted of a 20-second news segment in one of the following foreign languages: Creole, Macedonian, Marathi, Persian, Swahili and Ukranian. Each music block consisted of a 20-second segment of instrumental music from one of the following pieces: Bach Sinfonia No. 5, Smokey by Mountain, Bamboula by L.M Gottschalk, Spagnoletta Nuova by Fabritio Caroso, Kuhlau: Fantaisie for Solo Flute in D major (Op. 38, No. 3), and a violin rendition of the country song “When the right one comes along”.

A fixation cross was presented on the screen for the entire duration of the task. To ensure that subjects were attending to the auditory presentation, target trials (guitar plucks) occurred randomly about once a minute during the task. The participants were instructed to press a button with their right index finger every time a target trial was presented. Sounds were presented through an MRI-compatible Avotec Conformal Headset.

### MRS Session

MR Spectroscopy data was collected using the same scanner on a different day. During this second session, we first collected T1-weighted structural images using the same parameters as in the fMRI session. MRS data were acquired using a MEGA-PRESS sequence with the following parameters: TE=68ms (TE1=15ms, TE2=53ms), TR=1.8sec, 256 transients (128 ON interleaved with 128 OFF) of 4,096 data points; spectral width=5kHz, frequency selective editing pulses (14ms) applied at 1.9ppm (ON) and 7.46 ppm (OFF); total scan time about 8.5 minutes per voxel.

MRS data were collected from two 3cm x 3cm x 3cm voxels placed in the left and right auditory cortex (Figure 1), left and right ventrovisual cortex and left and right sensorimotor cortex (Figure S1). In order to ensure subject-level specificity, auditory voxels were placed to overlap maximally with each participant’s own functional activation maps (using a contrast of Speech + Music vs. Fixation) obtained from the fMRI run described previously.

**Fig 1.**
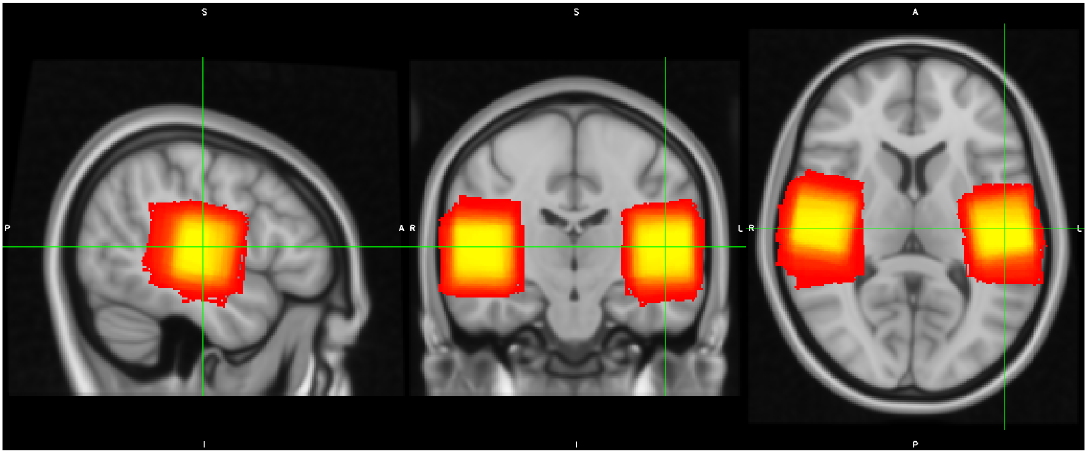
MRS voxel placement in the auditory cortex. The color indicates the amount of overlap in the voxel placement across participants (yellow represents maximum overlap while red represents less overlap).

### Quantification of GABA levels

We used the Gannet 3.0 MATLAB toolbox to estimate GABA levels in each of the two (left and right auditory) MRS voxels. The time domain data was frequency- and phase-corrected using spectral registration. It was filtered with 3-Hz exponential line broadening and zero-filled by a factor of 16. GABA levels were computed by fitting a Gaussian model to the 3-ppm peak in the difference spectrum and quantified relative to water (fit with a Gaussian-Lorentzian model) in institutional units (Figure S2). This editing scheme results in significant excitation of coedited macromolecule (MM) signal, that have been reported to contribute approximately 45% to the edited signal at 3-ppm. Thus, we report all GABA values as GABA+ (i.e., GABA + MM) in the present study. There are substantial differences in the relaxation constants and water visibility between WM, GM and CSF. To account for these differences, a binary mask of the MRS voxels was created using Gannet’s integrated voxel-to-image co-registration. Next, segmentation of the anatomical image was performed using the Segment function in SPM12 and the voxel fractions containing CSF, GM and WM were computed. From this procedure, a tissue-corrected GABA+ value was calculated for each participant. Since, the signal in GM and WM have different strengths an alpha tissue-corrected (fully corrected) GABA+ value was also computed for each participant.

### fMRl Data Preprocessing

fMRI data were k-space despiked, reconstructed, and corrected for heart beat and breathing using the RETROICOR algorithm. The initial five volumes were deleted and the data were then slice time corrected using the spm_slice_timing function from SPM8. Motion correction was performed using the Freesurfer FSFAST processing stream. Freesurfer was used to resample the data into two-dimensional cortical surfaces (one for the left hemisphere and one for the right hemisphere) based on a white/gray matter segmentation of each subject’s own high-resolution structural image computed using Freesurfer’s recon-all function. The data were then spatially smoothed within each cortical surface using a 5-mm two-dimensional smoothing kernel.

### ROI Selection

Because this was an auditory task we restricted our analysis to an anatomical mask containing the bilateral superior temporal gyrus, bank of the superior temporal sulcus, transverse temporal gyrus and supramarginal gyrus using cortical parcellation labels generated by FreeSurfer based on the Desikan-Killiany Atlas (aparc.annot). The resulting mask contained more than 37,000 vertices on the cortical surface (Figure 2). We obtained grey-matter thickness, volume and surface area estimates within this mask.

**Fig 2.**
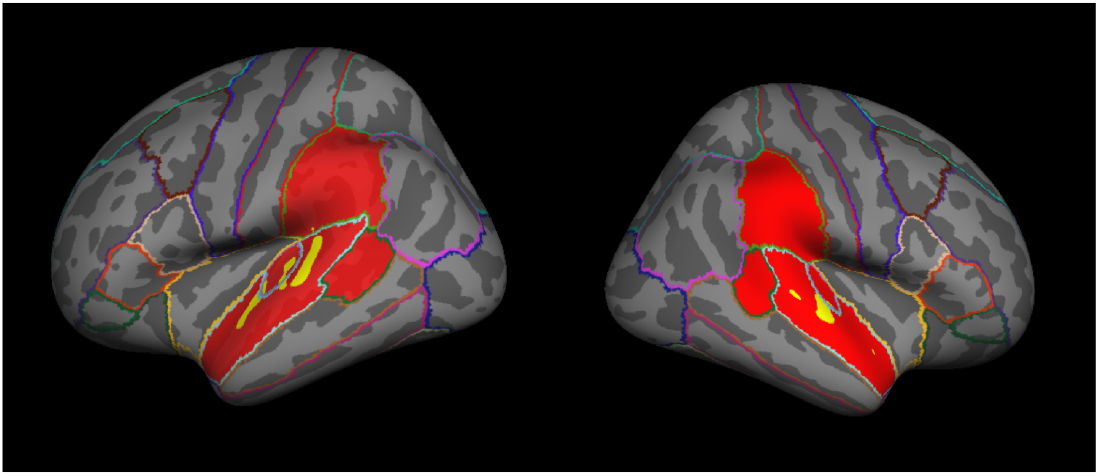
Participant-specific example of structural (in red) and functional (in yellow) masks. The functional mask was based on the 1400 most activated vertices from the music vs. fixation contrast (Fig. S3a) and the foreign speech vs. fixation contrast (Fig. S3b) under the constraint that an equal number of vertices were included from each contrast.

In order to ensure that only subject-specific, task-relevant vertices were analyzed, we then created a functional mask for each subject. Neural activation was estimated using a General Linear Model, fit with two box-car regressors (music vs. fixation and speech vs. fixation), convolved with a standard hemodynamic function. Beta values for each of the two regressors were obtained at each vertex. The functional mask was generated by selecting the most active vertices from both conditions in an alternating order (e.g., the most highly activated vertex for the music vs. fixation contrast, then the most highly activated vertex for the speech vs. fixation contrast, then the next most activated vertex for the music vs. fixation contrast, etc.). If the next most active vertex for a contrast had already been included in the functional mask, then the next most active voxel that had not already been included in the functional mask was added. This approach ensured that both conditions were equally represented in the functional mask. The functional mask selection was blind to whether the chosen vertex was selective for one condition or was activated by both conditions (Figure 2).

Using a speech + music vs. fixation contrast, we calculated the total number of vertices across both hemispheres that were activated (p<0.001, uncorrected) during auditory perception for each subject. 95% of the subjects had greater than 1400 such vertices, so we chose an ROI-size of 1400 vertices as our default functional mask size. We also varied the ROI-size from small (1000 vertices) to very large (the entire anatomical mask) to ensure that any observed effects on neural distinctiveness did not depend on the size of the ROI.

### Neural activation

In order to generate multiple independent activation patterns for use in multivoxel pattern analysis (MVPA), we then fit another General Linear Model that included separate box-car regressors for each of the 12 task-blocks (6 music and 6 speech), convolved with a standard hemodynamic function. Fitting the model produced beta values at each vertex separately for each of the 12 blocks. Neural distinctiveness was computed using these beta values (activation maps) as described below.

### SVM-based calculation of distinctiveness

Machine learning algorithms, particularly linear-SVMs (support vector machines) provide a measure of the distinctiveness of different patterns of neural activation. A linear SVM finds a hyperplane that maximally separates neural patterns into two different categories provided during training. Then this trained SVM is used to classify untrained neural patterns into one of the two categories. We used a leave-one-out cross-validation approach, in which the classifier was trained to fit 11 of the 12 activation maps (6 music and 6 speech) within the functional ROI and then was tested on the remaining activation map. This process was repeated leaving out each of 12 different activation maps and the average classification accuracy was used as a measure of neural distinctiveness. This measure of classification accuracy can only take on one of 13 discrete values (100%, 92%, 83%, 75%, 66%, etc.) corresponding to the number of the 12 activation maps that were classified correctly (plus 0% accuracy). Classification accuracy of 50% is chance.

### Correlation-based calculation of distinctiveness

We also used a correlation-based approach that produces a more continuous measure of neural distinctiveness and that avoids ceiling effects (11, 31). For each subject, correlations between the activation maps for all pairs of blocks of the same type were computed within the functional ROI (e.g., music block 1 with music block 2, music block 3 with music block 6, speech block1 with speech block4, etc.). These correlations were then averaged to produce a within-category correlation value. Likewise, correlations between activation maps for all pairs of blocks of different types were computed (e.g., music block 1 with speech block 2, music block 3 with speech block 6, speech block1 with music block4, etc.). These correlations were then averaged to produce a between-category correlation value. Neural distinctiveness was then defined as the difference between the average within-category correlation and average between-category correlation. This measure has a theoretical range of 2 to −2. This multivariate analysis reveals fine-grained differences in the distinctiveness of activation patterns rather than differences in the average activation between the two categories as a univariate method would.

## Results

### Neural Distinctiveness and Aging

Neural distinctiveness as measured by SVM classifier accuracy (Figure 3a) was significantly lower in older adults (mean = 85.9%) compared to young adults (mean = 96.3%), (t (41) = −3.06, p = 0.004). Likewise, when neural distinctiveness was computed based on pattern similarity/dissimilarity using the difference between within-category and between-category correlations (Figure 3b), older adults exhibited less distinctive activation patterns (mean = 0.27) than did young adults (mean = 0.39) (t (41) = −2.04, p = 0.047). In other words, using both measures the activation patterns for music and speech were more similar or confusable in older adults than younger adults.

**Fig 3.**
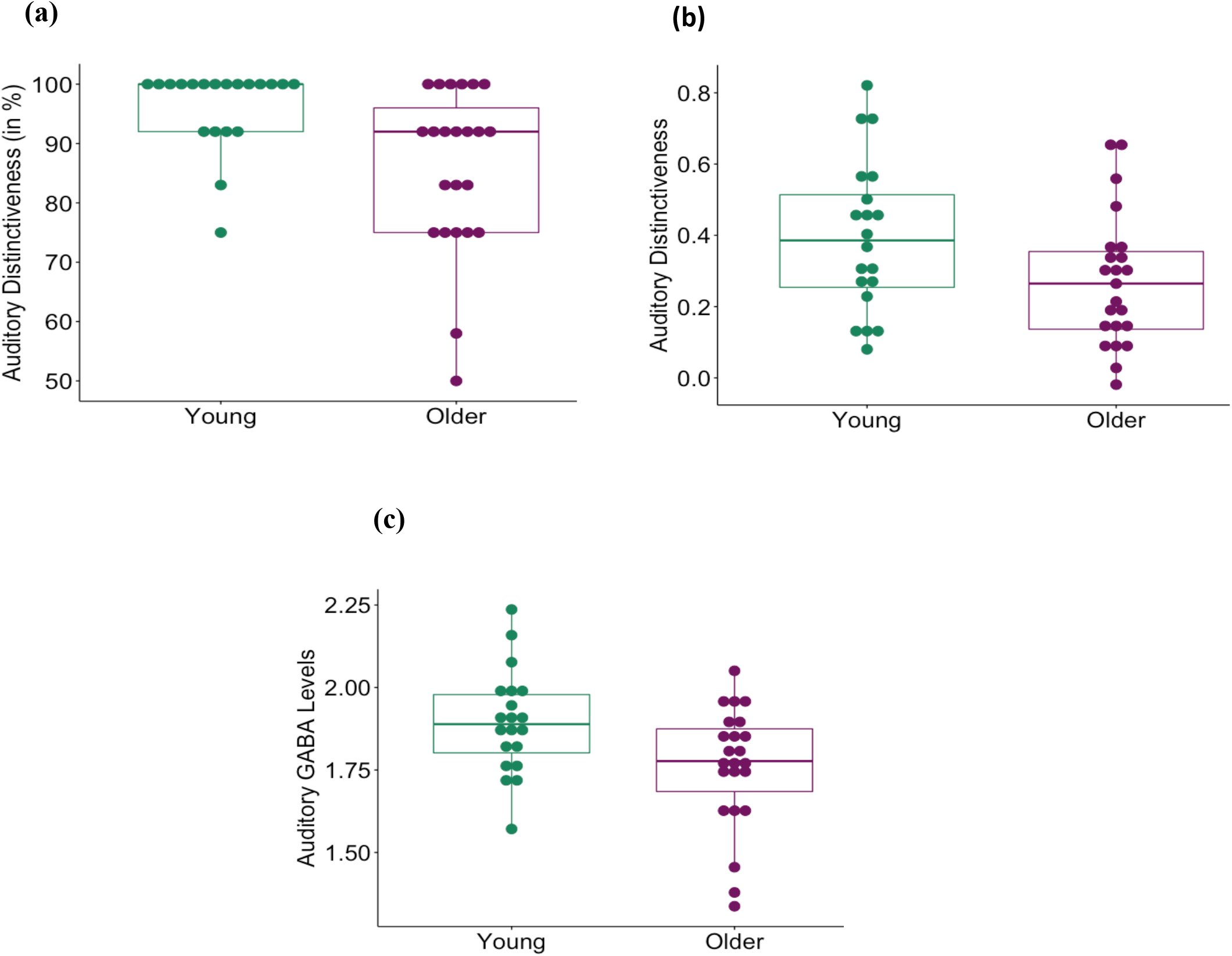
(a) Neural distinctiveness based on the accuracy of an SVM classifier in distinguishing activation patterns evoked by music from those evoked by foreign speech. Distinctiveness was significantly lower in older adults (in purple) than young adults (in green) (t (41) = −3.065, p = 0.004). (b) Neural distinctiveness based on the difference between within-condition similarity and between-condition similarity. Distinctiveness was again significantly lower in older adults (in purple) than young adults (in green) (t (41) = −2.04, p = 0.047). (c) Raw GABA+/Water levels in the auditory cortex estimated by MRS. GABA+ levels were significantly lower in older adults (in purple) than young adults (in green) (t (41) = −2.6, p = 0.01).

With 12 activation patterns to classify, the SVM-based measure of distinctiveness can only take on 13 different values. It is also prone to ceiling effects (e.g., the classifier was 100% accurate in classifying the activation patterns for 20 of the 43 participants). In contrast, the correlation-based measure can take on any real value between −2 and 2 and is much less susceptible to ceiling effects. The two measures were also significantly correlated (r (41) = 0.42, p = 0.004). We therefore used the correlation-based measure for subsequent analyses.

There was no significant difference (t (41) = −0.64, p = 0.53) between the number of activated vertices (p<0.001, uncorrected) within the anatomical mask for young (mean = 7155) and older adults (mean = 6491). Furthermore, there was no significant difference in the mean (t (41) = −1.26, p=0.22) or peak (t (41) = −0.71, p=0.48) activation level between the two age groups. Differences in distinctiveness between the age-groups were therefore not driven by differential activation levels between the two age groups, but rather by age differences in the similarity/dissimilarity of neural activation patterns elicited by music and speech.

In order to ensure that the effect of aging on neural distinctiveness was not due to the selection of a particular ROI size, we computed a pairwise t-test at every ROI size and found that distinctiveness declined with age independent of ROI size selection (the effect was only marginally significant at the smallest ROI size) (Figure 4, Table S1). However, as ROI-size increased, the average distinctiveness values declined suggesting that the larger ROIs included task-irrelevant vertices that added noise to the distinctiveness measure.

**Fig 4.**
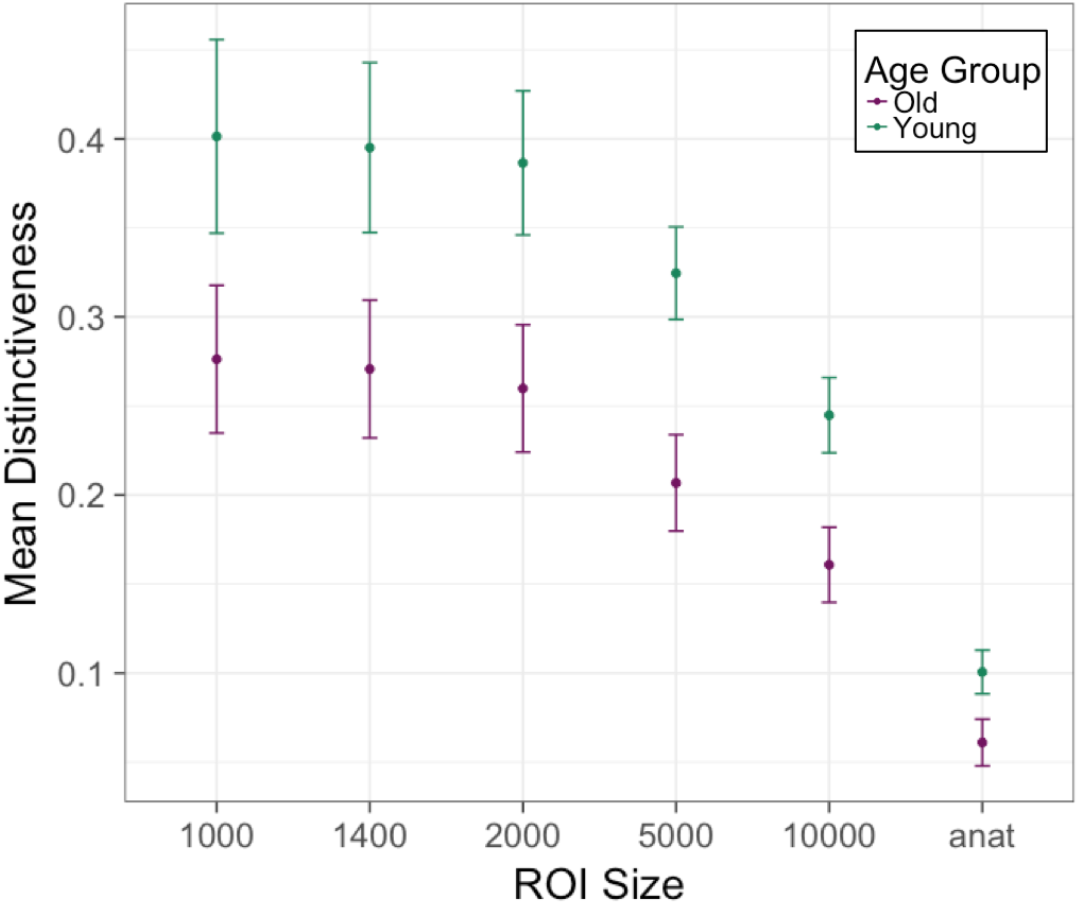
Neural distinctiveness in the two age groups as a function of ROI size. Distinctiveness was significantly lower in older adults (in purple) than younger adults (in green) for most ROI sizes (also see Table 1). The vertical axis is mean distinctiveness (measured as within-between difference) with standard error bars. The horizontal axis is the ROI size (in number of vertices; anat refers to the entire anatomical mask of approximately 37000 vertices).

The observed age-related decline in neural distinctiveness could be due to changes in the ear, rather than changes in the brain. That is, peripheral changes in the ear that reduce auditory sensitivity could lead to reduced neural distinctiveness, even if there were no age-related changes in auditory cortex itself. To explore this issue, we analyzed neural distinctiveness after controlling for individual participants’ pure-tone threshold. We still found that neural distinctiveness was significantly lower in the older vs. younger participants (r (41) = −0.30, p = 0.048). Thus, although peripheral changes may contribute to age-related declines in auditory neural distinctiveness, they do not completely explain them.

We also examined age-related changes in grey-matter thickness and surface area. Older adults exhibited significantly thinner grey-matter (t (37.4) =-6.82, p= 4.7e-08) and reduced surface area (t (39.6) =-3.49, p=0.001) within the anatomically defined mask. Neural distinctiveness was still significantly lower in the older adults even after controlling for changes in grey-matter thickness (r (41) =-0.32, p=0.037), but not after controlling for surface area (r (41) = −0.17, p= 0.28). These results indicate that changes in neural distinctiveness might be at least partially due to anatomical changes that accompany aging.

### GABA+ Levels and Aging

Raw GABA+ levels were significantly lower in the auditory cortex in older adults (mean = 1.75) than in young adults (mean = 1.89) (t (40.9) =-2.78, p=0.008) (Figure 3c).

We also used an ANCOVA to investigate whether there were systematic differences between GABA levels across hemisphere and if this effect interacted with age. There was a significant main effect of age on GABA+ independent of hemisphere (F (1,41) = 7.5, p=0.009) but no main effect of hemisphere (F (1,41) = 0.008, p=0.93). There was also no significant interaction between hemisphere and age (F (1,41) = 0.19, p=0.66). Because there were no significant differences between the GABA+ estimates in the two hemispheres and because the two estimates were significantly correlated (r (41) = 0.52, p=0.0003), we averaged the GABA+ estimates from each hemisphere for further analysis.

Gannet also provides GABA+ estimates that account for differences in relaxation times of GABA and water based on tissue composition (i.e., the fraction of grey matter, white matter and cerebrospinal (CSF) volume within each voxel). GABA+ levels were lower in older adults (mean = 1.99) than young adults (mean = 2.18) even after these corrections (t (40.5) = −3.24, p = 0.002). Gannet also provides GABA+ estimates that account for difference in GABA signal strength in white matter compared to grey matter. Age did not have a significant main effect on these fully corrected GABA+ estimates (t (40.45) = 0.26, p = 0.8). These results indicate that accounting for structural changes with age like tissue composition can explain age differences in GABA+ estimates in auditory cortex.

### GABA and Distinctiveness

Average raw GABA+ levels in the auditory cortex were positively correlated with neural distinctiveness in the older adults (r (21) = 0.54, p = 0.008) (Figure 5), but not the younger adults (r (18) = −0.18, p = 0.45) (Figure S4). This GABA-distinctiveness relationship was also region-specific: neither ventrovisual GABA (r (21) = 0.25, p = 0.25) nor sensorimotor GABA (r (21) = 0.19, p = 0.38) were significantly correlated with auditory distinctiveness in the older adults. This relationship between auditory GABA and auditory distinctiveness was also still significant after controlling for age within the older adults (r (21) = 0.54, p = 0.009).

**Fig 5.**
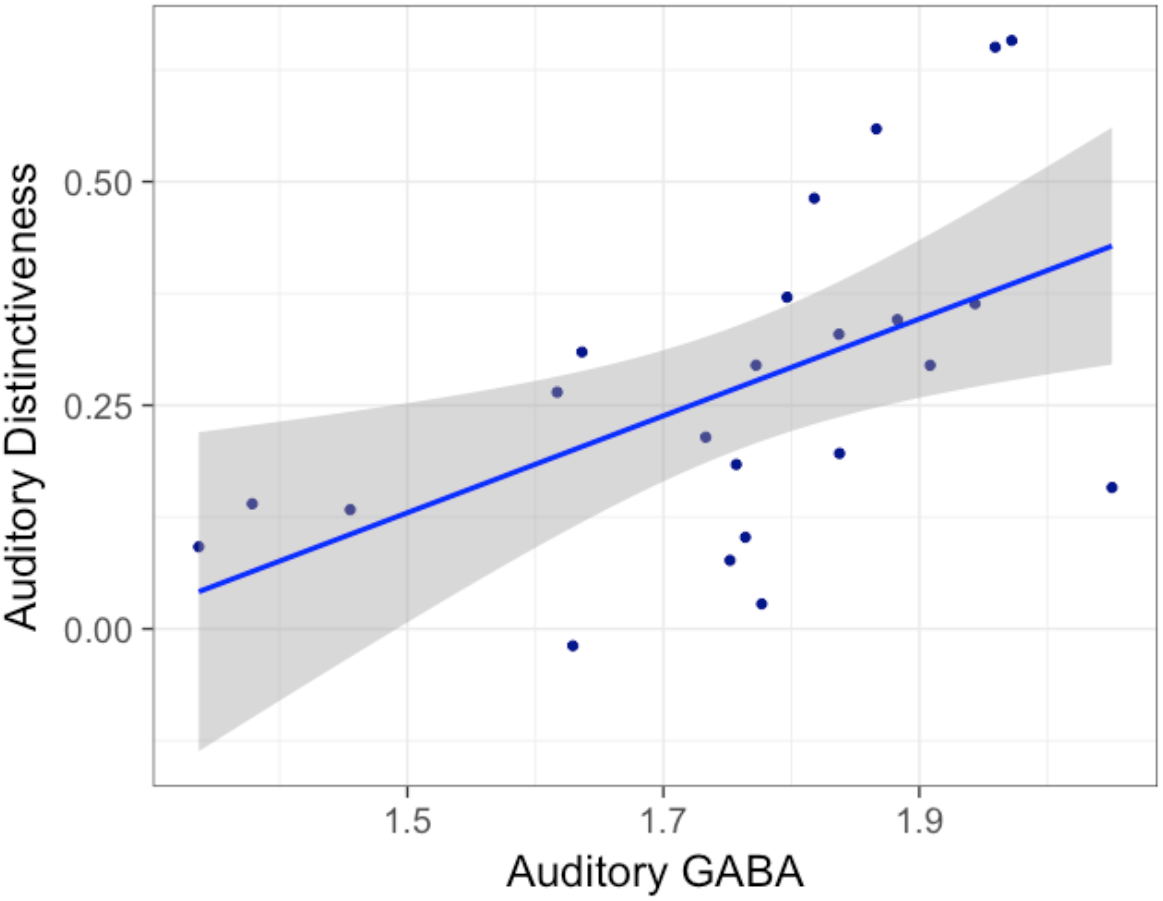
The relationship between raw auditory GABA+ levels and auditory neural distinctiveness in older adults. Individual differences in GABA+ were significantly correlated with individual differences in neural distinctiveness in older adults. (r (21) = 0.54, p = 0.008)

We also examined fully alpha, tissue-corrected GABA+ estimates to investigate whether the relationship between neural distinctiveness and GABA was driven by structural changes that accompany aging. We also included age and gray matter volume as nuisance covariates. The main effect of fully-corrected GABA+ on distinctiveness was still significant (F (1,18) = 4.44, p = 0.049), but the effect of age (F (1,18) = 0.01, p=0.92) and grey matter volume (F (1,18) = 0.60, p = 0.45) were not. Thus, the relationship between GABA and neural distinctiveness cannot be fully explained by age-related changes in grey-matter volume.

## Discussion

The age-related neural dedifferentiation hypothesis posits that the neural representations of different stimuli become less distinct with age (32). Most of the previous evidence for this hypothesis in humans has come from studies of visual cortex. In the present study, we showed that neural distinctiveness also declines with age in the auditory cortex, extending the scope of neural dedifferentiation theory. This age-related decline in distinctiveness was independent of ROI-size, was present after controlling for peripheral hearing performance (pure-tone threshold), and was still present after controlling for grey matter thickness.

We also examined GABA levels in the auditory cortex and the relationship between GABA and distinctiveness. Consistent with previous animal research, we found that GABA levels decline with age in the auditory cortex and showed for the first time that individual differences in GABA levels predict individual differences in neural distinctiveness.

### Age-related dedifferentiation

Previous research in animals, provides direct evidence for age-related decline in neural selectivity or distinctiveness in single neurons (23, 33, 34). Neuroimaging studies in humans have also found that neural patterns of activation in large cortical regions become less distinct with age in ventral visual cortex while viewing faces vs. houses (9, 35, 36); in motor cortex during left vs. right finger tapping (8), in hippocampus during memory retrieval of different items (37) and in posterior medial cortex for different emotion regulation strategies (38). Our study contributes to this growing body of literature by showing that age-related dedifferentiation extends to auditory cortex.

A natural question is whether the observed declines in neural distinctiveness in auditory cortex are due to age-related changes in the peripheral auditory system, i.e. the ear, or whether they reflect more central changes in the cortex. Aging is accompanied by several changes in the ear, including the loss of hair cells, dysfunction of the stria vascularis, and stiffening of the basilar membrane (39). Such changes in the peripheral auditory system could result in a noisier auditory input. And noisier information could plausibly produce less distinctive cortical representations, even if central auditory processing in the cortex itself has not changed dramatically. However, we observed age-related dedifferentiation in our sample even after controlling for pure-tone thresholds, suggesting that age-related changes in the auditory cortex itself may also be playing a role in auditory dedifferentiation. Consistent with this hypothesis, age-related declines of central auditory processing often appear even after accounting for peripheral hearing loss (40–43).

### Age-related decline in GABA levels

Several animal studies have reported that levels of the inhibitory neurotransmitter GABA decline with age in the auditory system. For example, GABA levels decline in the inferior colliculus (25, 26, 39) and auditory cortex (44) of aging rats, as well as in the cochlea of aging mice (45). GABA receptor binding and subunit levels also decline with age (27, 28).

Only a few human studies have investigated age-related changes in auditory GABA levels, and the results are mixed. Profant et al. (2015) did not observe a significant effect of age on GABA levels in auditory cortex. In contrast, Chen et al. (2013) did report a significant decline in GABA levels: in the right (but not the left) hemisphere before pure tone stimulation, and in both hemispheres after stimulation. Likewise, Gao et al. (2015) reported that older adults suffering from age-related hearing loss exhibited lower GABA levels in auditory cortex compared to other older adults. Consistent with these results, our study provides further evidence that auditory GABA levels decline significantly with age in older adults compared to younger adults. However, fully corrected GABA estimates did not show an age-related decline. This suggests that observed declines in GABA levels with age may be mediated by changes in tissue composition with aging.

The observed age-related declines in GABA are also consistent with the view that some age-related behavioral impairments may reflect an underlying deficit in inhibition (46, 47). These theories suggest that as a result of impaired inhibitory function, older adults have greater difficulty preventing irrelevant information from gaining access to attention than young adults. Thus, older adults may be more susceptible to distraction and more likely to choose a non-optimal response. Since GABA is the brain’s major inhibitory neurotransmitter, age-related reductions in GABA could naturally explain the observed inhibitory deficit.

### Auditory GABA decline predicts dedifferentiation

Leventhal et al. (2003) showed that the neural selectivity of orientation-specific cells in visual cortex declines with age. They also showed that the selectivity of individual neurons can be experimentally manipulated by the application of GABA, a GABA agonist, or a GABA antagonist. Specifically, visual neurons in older macaques that were not orientation-selective became selective after the application of GABA or the GABA agonist muscimol. Conversely, visual neurons in young macaques that were orientation-selective, became non-selective after the application of the GABA antagonist bicuculline. Together these results demonstrate that changes in GABA activity can cause changes in neural selectivity, at least in individual neurons in visual cortex.

Researchers have reported similar findings in the auditory system. For example, the application of a GABA antagonist reduces the selectivity of cells to sinusoidally amplitude modulated (SAM) stimuli in the inferior colliculus of rats (19), as well as the rate and direction selectivity of cells to FM sweeps in the auditory system of bats (48).

Obviously, the selectivity of individual receptive fields might be quite different from the selectivity of the large-scale neural representations that can be measured using fMRI in humans. Nevertheless, age-related declines in GABA could plausibly influence both and so we decided to test whether individual differences in GABA were associated with individual differences in neural distinctiveness. And the results confirmed the prediction. Older participants had significantly greater neural distinctiveness than did older adults with lower GABA levels, even after controlling for age and tissue composition differences. These results are consistent with the hypothesis that age-related declines in GABA contribute to age-related neural dedifferentiation.

Furthermore, this relationship was region-specific: GABA estimates in ventrovisual and somatosensory cortex were not significantly associated with auditory distinctiveness. These results suggest that the observed GABA-distinctiveness relationship is probably not due to some confounding effect (increased variance with age, vascular changes with age) that would be present throughout the brain.

Animal research has shown a direct association between decline in auditory neural selectivity and age-related hearing loss (33, 49). If GABA levels influence neural distinctiveness, as our results suggest, then pharmacological treatments that target GABA could be a promising avenue for clinical research aimed at mitigating age-related hearing impairments.

### Limitations

A key limitation of the current study is that it is correlational. We therefore cannot conclude that age-related changes in GABA cause changes in neural distinctiveness, only that they are related. Another limitation is that MRS estimates of GABA do not measure GABA activity, but GABA volume. Nor do they distinguish between intracellular and extracelluar GABA. Furthermore, the measurements are made in a fairly large volume that does not overlap exactly with the functional region of interest. These shortcomings should presumably make it harder to observe relationships between auditory GABA and auditory distinctiveness, so the fact that we did find a significant relationship suggests that the relationship may be fairly strong.

## Conclusions

In sum, our findings show that sensory neural dedifferentiation is not limited to ventral visual cortex but extends to the auditory cortex. Furthermore, they demonstrate that GABA levels in auditory cortex decline with age and that individual differences in GABA predict individual differences in neural distinctiveness. Together these findings are consistent with the hypothesis that age-related declines in GABA contribute to age-related declines in neural distinctiveness.

## Acknowledgements

This work was supported by a grant from the National Institutes of Health to TAP (RA01AG050523). The authors would like to thank Dr. Cindy Lustig for valuable comments on previous drafts.

## References

1. Harada CN, Natelson Love MC, Triebel K (2013) Normal Cognitive Aging. Clin Geriatr Med 29(4):737–752.

2. Park DC, et al. (2002) Models of visuospatial and verbal memory across the adult life span. Psychol Aging 17(2):299–320.

3. Salthouse TA (1996) The processing-speed theory of adult age differences in cognition. Psychol Rev 103(3):403–428.

4. Fortunato S, et al. (2016) A review of new insights on the association between hearing loss and cognitive decline in ageing. Acta Otorhinolaryngol Ital 36(3):155–166.

5. Christensen H, et al. (1999) An analysis of diversity in the cognitive performance of elderly community dwellers: individual differences in change scores as a function of age. Psychol Aging 14(3):365–379.

6. Hultsch DF, MacDonald SWS, Dixon RA (2002) Variability in reaction time performance of younger and older adults. J Gerontol B Psychol Sci Soc Sci 57(2):P101–115.

7. Wilson RS, et al. (2002) Individual differences in rates of change in cognitive abilities of older persons. Psychol Aging 17(2):179–193.

8. Carp J, Park J, Hebrank A, Park DC, Polk TA (2011) Age-related neural dedifferentiation in the motor system. PLoS ONE 6(12). doi:10.1371/journal.pone.0029411.

9. Park DC, et al. (2004) Aging reduces neural specialization in ventral visual cortex. Proc Natl Acad Sci U S A 101(35):13091–13095.

10. Voss MW, et al. (2008) Dedifferentiation in the visual cortex: an fMRI investigation of individual differences in older adults. Brain Res 1244:121–131.

11. Park J, Carp J, Hebrank A, Park DC, Polk TA (2010) Neural specificity predicts fluid processing ability in older adults. J Neurosci Off J Soc Neurosci 30(27):9253–9259.

12. Chalavi S, et al. (2018) The neurochemical basis of the contextual interference effect. Neurobiol Aging 66:85–96.

13. Hermans L, et al. (2018) Brain GABA levels are associated with inhibitory control deficits in older adults. J Neurosci Off J Soc Neurosci. doi:10.1523/JNEUROSCI.0760-18.2018.

14. Simmonite M, et al. (2018) Age-related declines in occipital GABA are associated with reduced fluid processing ability. Acad Radiol. doi:10.1016/j.acra.2018.07.024.

15. Gao F, et al. (2013) Edited magnetic resonance spectroscopy detects an age-related decline in brain GABA levels. NeuroImage 78:75–82.

16. Cassady, K., Gagnon, H., Lalwani, P., Simmonite, M., Foerster, B., Park, D., Peltier, S.J., Petrou, M., Taylor, S., Weissman, D.H., Seidler, R.D., & Polk, T.A. (in press). Sensorimotor network segregation declines with age and is linked to GABA and sensorimotor performance. NeuroImage.

17. Porges EC, et al. (2017) Frontal gamma-aminobutyric acid concentrations are associated with cognitive performance in older adults. Biol Psychiatry Cogn Neurosci Neuroimaging 2(1):38–44.

18. Leventhal AG, Wang Y, Pu M, Zhou Y, Ma Y (2003) GABA and its agonists improved visual cortical function in senescent monkeys. Science 300(5620):812–815.

19. Caspary DM, Palombi PS, Hughes LF (2002) GABAergic inputs shape responses to amplitude modulated stimuli in the inferior colliculus. Hear Res 168(1–2):163–173.

20. Chen X, et al. (2013) Age-associated reduction of asymmetry in human central auditory function: A 1H-magnetic resonance spectroscopy study. Neural Plast 2013. doi:10.1155/2013/735290.

21. Profant O, et al. (2015) Functional changes in the human auditory cortex in ageing. PLoS ONE 10(3). doi:10.1371/journal.pone.0116692.

22. Villers-SidaniE de, et al. (2010) Recovery of functional and structural age-related changes in the rat primary auditory cortex with operant training. Proc Natl Acad Sci 107(31):13900–13905.

23. Juarez-Salinas DL, Engle JR, Navarro XO, Recanzone GH (2010) Hierarchical and serial processing in the spatial auditory cortical pathway is degraded by natural aging. J Neurosci 30(44):14795–14804.

24. Leong U-C, Barsz K, Allen PD, Walton JP (2011) Neural correlates of age-related declines in frequency selectivity in the auditory midbrain. Neurobiol Aging 32(1):168–178.

25. Caspary DM, Raza A, Lawhorn Armour BA, Pippin J, Arnerić SP (1990) Immunocytochemical and neurochemical evidence for age-related loss of GABA in the inferior colliculus: implications for neural presbycusis. J Neurosci Off J Soc Neurosci 10(7):2363–2372.

26. Gutiérrez A, Khan ZU, Morris SJ, De Blas AL (1994) Age-related decrease of GABAA receptor subunits and glutamic acid decarboxylase in the rat inferior colliculus. J Neurosci Off J Soc Neurosci 14(12):7469–7477.

27. Caspary DM, Hughes LF, Ling LL (2013) Age-related GABAA receptor changes in rat auditory cortex. Neurobiol Aging 34(5):1486–1496.

28. Milbrandt JC, Albin RL, Caspary DM (1994) Age-related decrease in GABAB receptor binding in the Fischer 344 rat inferior colliculus. Neurobiol Aging 15(6):699–703.

29. Carson N, Leach L, Murphy KJ (2018) A re-examination of Montreal Cognitive Assessment (MoCA) cutoff scores. Int J Geriatr Psychiatry 33(2):379–388.

30. Gagnon H, et al. (2018) Michigan Neural Distinctiveness (MiND) project: Investigating the scope, causes, and consequences of age-related neural dedifferentiation. bioRxiv:466516.

31. Haxby JV, et al. (2001) Distributed and overlapping representations of faces and objects in ventral temporal cortex. Science 293(5539):2425–2430.

32. Li SC, Lindenberger U, Sikström S (2001) Aging cognition: From neuromodulation to representation. Trends Cogn Sci 5(11):479–486.

33. Khouri L, Lesica NA, Grothe B (2011) Impaired auditory temporal selectivity in the inferior colliculus of aged Mongolian gerbils. J Neurosci Off J Soc Neurosci 31(27):9958–9970.

34. Schmolesky MT, Wang Y, Pu M, Leventhal AG (2000) Degradation of stimulus selectivity of visual cortical cells in senescent rhesus monkeys. Nat Neurosci 3(4):384–390.

35. Goh JO, Suzuki A, Park DC (2010) Reduced neural selectivity increases fMRI adaptation with age during face discrimination. NeuroImage 51(1):336–344.

36. Goh JOS (2011) Functional dedifferentiation and altered connectivity in older adults: Neural accounts of cognitive aging. Aging Dis 2(1):30–48.

37. Giovanello KS, Schacter DL (2012) Reduced specificity of hippocampal and posterior ventrolateral prefrontal activity during relational retrieval in normal aging. J Cogn Neurosci 24(1). doi:10.1162/jocn_a_00113.

38. Martins B, Ponzio A, Velasco R, Kaplan J, Mather M (2015) Dedifferentiation of emotion regulation strategies in the aging brain. Soc Cogn Affect Neurosci 10(6):840–847.

39. Ouda L, Syka J (2012) Immunocytochemical profiles of inferior colliculus neurons in the rat and their changes with aging. Front Neural Circuits 6:68.

40. Brant LJ, Fozard JL (1990) Age changes in pure-tone hearing thresholds in a longitudinal study of normal human aging. J Acoust Soc Am 88(2):813–820.

41. Dubno JR, Dirks DD, Morgan DE (1984) Effects of age and mild hearing loss on speech recognition in noise. J Acoust Soc Am 76(1):87–96.

42. Duquesnoy AJ (1983) The intelligibility of sentences in quiet and in noise in aged listeners. J Acoust Soc Am 74(4):1136–1144.

43. Souza PE, Boike KT, Witherell K, Tremblay K (2007) Prediction of speech recognition from audibility in older listeners with hearing loss: Effects of age, amplification, and background noise. doi:info:doi/10.3766/jaaa.18.1.5.

44. Ling LL, Hughes LF, Caspary DM (2005) Age-related loss of the GABA synthetic enzyme glutamic acid decarboxylase in rat primary auditory cortex. Neuroscience 132(4):1103–1113.

45. Tang X, et al. (2014) Age-related hearing loss: GABA, nicotinic acetylcholine and NMDA receptor expression changes in spiral ganglion neurons of the mouse. Neuroscience 259:184–193.

46. Hasher L, Zacks RT (1988) Working memory, comprehension, and aging: A review and a new view. Psychology of Learning and Motivation, ed Bower GH (Academic Press), pp 193–225.

47. Lustig C, Hasher L, Zacks RT (2007) Inhibitory deficit theory: Recent developments in a “new view.” Inhibition in Cognition (American Psychological Association, Washington, DC, US), pp 145–162.

48. Razak KA, Fuzessery ZM (2009) GABA shapes selectivity for the rate and direction of frequency-modulated sweeps in the auditory cortex. J Neurophysiol 102(3):1366–1378.

49. Trujillo M, Razak KA (2013) Altered cortical spectrotemporal processing with age-related hearing loss. J Neurophysiol 110(12):2873–2886.

50. Turner JG, Hughes LF, Caspary DM (2005) Affects of aging on receptive fields in rat primary auditory cortex layer V neurons. J Neurophysiol 94(4):2738–2747.

